# A Model for Genome-First Care: Returning Secondary Genomic Findings to Participants and Their Healthcare Providers in a Large Research Cohort

**DOI:** 10.1101/166975

**Authors:** Marci L. B. Schwartz, Cara Zayac McCormick, Amanda L. Lazzeri, D’Andra M. Lindbuchler, Miranda L. G. Hallquist, Kandamurugu Manickam, Adam H. Buchanan, Alanna Kulchak Rahm, Monica A. Giovanni, Lauren Frisbie, Carroll N. Flansburg, F. Daniel Davis, Amy C. Sturm, Christine Nicastro, Matthew S. Lebo, Heather Mason-Suares, Lisa Marie Mahanta, David J. Carey, Janet L. Williams, Marc S. Williams, David H. Ledbetter, W. Andrew Faucett, Michael F. Murray

## Abstract

**Background:** Research cohorts with linked genomic data exist, or are being developed, at many research centers. Within any such “sequenced cohort” of more than 100 participants, it is likely that there are participants with previously undisclosed risk for life-threatening monogenic diseases that could be identified with targeted analysis of their existing data. Identification of such disease-associated findings are not usually primary to the enrollment research goals. At Geisinger Health System, MyCode^®^ Community Health Initiative (MyCode) participants represent one such large sequenced cohort. Since 2013, MyCode participants in discovery research have been consented for secondary analysis of their existing research genomic sequences to allow delivery of medically actionable findings to them and their healthcare providers. This return of genomic results program was developed to manage an anticipated 3.5% of MyCode participants who will receive clinically confirmed genomic variants from an approved gene list out of more than 150,000 total participants. Risk-associated DNA sequences alone without any clinical parameter, prompt “genome-first” follow-up encounters.

**Methods:** This article describes our process for generating clinical grade results from research-based genomic sequencing data, delivering results to patients and their providers, facilitating targeted clinical evaluations of patients and promoting cascade testing of at-risk relatives. We also summarize our early data about the results generated during this process and our ability to contact patients and their providers to disclose the information.

**Results:** This process has been used to generate 343 results on 339 patients. 93% of patients with a result have been successfully contacted about their results as evidenced by direct interaction about their result with the research team or a healthcare provider. 222 healthcare providers have been notified of a result on one or more patient through this result delivery process.

**Conclusions:** Here we describe the existing GHS model to deliver genomic data into the electronic medical record and the clinical interactions that are prompted and supported. Elements of this genome-first care model can be applied in other healthcare settings and in national efforts, such as “All of Us”, that wish to establish programs for returning genomic results to research participants.

## BACKGROUND

Genomic sequencing is increasingly used in diagnostic and research applications due to decreasing costs and improvements in informatics analysis and public genomics databases. [1] This has led to the development of guidelines about the return of incidental or secondary findings. [2] The American College of Medical Genetics and Genomics (ACMG) proposed an initial list of 56 genes (subsequently revised to 59 genes), to be analyzed for variants known or expected to confer disease risk even when unrelated to the primary indication for clinical testing.[3] These guidelines only apply to clinical sequence data. Similar guidelines do not exist for sequences generated in research.[3, 4]

A large sequenced cohort has been established at Geisinger Health System (GHS) using biospecimens collected through an EHR-linked biobank, The MyCode^®^ Community Health Initiative (MyCode). [5] MyCode specimens are utilized for collaborative research with investigators at the Regeneron Genomics Center. [6] Over 150,000 Geisinger patients have consented to participate in MyCode with an enrollment goal of at least 250,000 participants. These participants are concentrated in the GHS service area which extends throughout central and northeastern Pennsylvania and southern New Jersey. Three key infrastructural components of MyCode enable the use of research-based sequence data for clinical care: 1) sequence data is linked with participants and their electronic health records (EHR), 2) participants have consented to receive results, and 3) biobank samples meet Clinical Laboratory Improvement Amendment (CLIA) standards and are available for clinical confirmatory genetic testing. An institutional decision was made to analyze genomic sequence data generated from the MyCode study to return medically actionable findings to participants.[7]

A return-of-results program was developed to identify genomic variants within the research data and to notify participants and their providers of this medically actionable information. This program represents an emerging application of precision health: incorporating a genome-first model where genomic data is used to identify individuals with highly penetrant expected pathogenic sequence variants which then trigger targeted screening, prevention, and interventions. This program does not use clinical disease or symptom development as the trigger for engaging genetic services. Rather it enables proactive care by informing participants of an elevated risk and engaging them in personalized screening and preventive management, ideally prior to disease development. Despite the clinical overlap, MyCode participation is not recommended to patients as a method for obtaining diagnostic genetic testing for individuals with a clinical indication, such as those with a personal or family history suggestive of a monogenic condition. Funding for this return-of-results program was obtained from institutional support, grants, and donations and whole exome sequence data was generated for research use through an industry partnership with Regeneron Pharmaceuticals.

An analysis of the research sequence data suggests that approximately 3.5% of MyCode participants will carry a pathogenic or likely pathogenic variant using the Geisinger 76 (G76) gene list [6] which is consistent with the range of 1.2%-6.2% reported in other studies. [8-10] Geisinger supports the Learning Healthcare System (LHS) framework which promotes the dual goal of applying existing clinical knowledge in new ways for the purpose of improving health outcomes and building a robust evidence-base for alternative practice models.[11] The MyCode return of results program is the first large scale example of a genomics-informed LHS.[12] The approach described here can be customized for any program that has existing sequence data linked to medical records to identify individuals for preemptive health interventions.

Organizing principles of the program include careful selection of medically actionable genes with potential for personal and population health impact, high-quality, stringent variant calling to minimize the false-positive rate, and a return process that supports patients, families, and clinicians with collaborative approaches to disclosure, evaluation, risk management, and cascade testing. These principles were important for this screening program because a positive result can lead to invasive risk management procedures for index patients and their family members. This paper describes the steps implemented to achieve the goals listed below:

1. Generate clinical grade results from research-based genomic sequencing data
2. Deliver results to healthcare providers utilizing the EHR
3. Communicate results to participants with explanation of the implications
4. Perform risk-specific targeted clinical evaluations
5. Facilitate cascade testing of at-risk relatives

## METHODS

### 1. Generate Clinical Grade Results from Research-based Genomic Sequencing Data

Several steps were implemented to generate a clinical result that meets a standard acceptable for use in clinical care and placement into the EHR. These steps include determination of a list of genes for analysis and return, sequence variant analysis and interpretation, verification of participant eligibility to receive a result, and confirmatory sequencing with subsequent reporting of positive findings (Figure 1).

**FIGURE 1:**
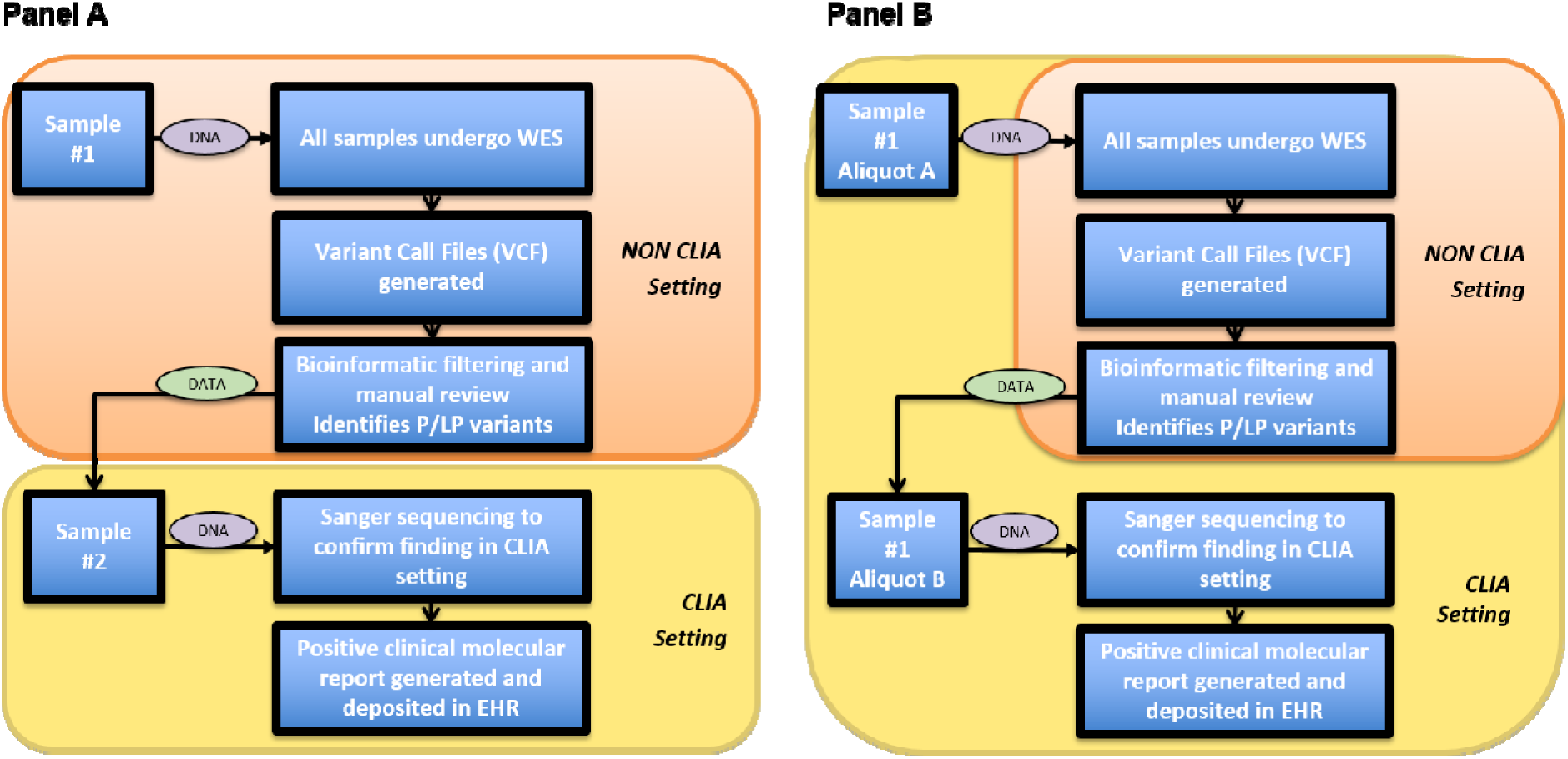
Figure 1 - Clinical Confirmation of Research Results. **Panel A** demonstrates the pathway for samples from participants recruited to MyCode prior to the biorepository’s CLIA certification in 2013. In such cases, it is necessary to collect a second sample for CLIA confirmation. **Panel B** demonstrates the pathway for samples from participants recruited to MyCode since the biorepository’s CLIA certification in 2014. In such cases a second aliquot (#1B) is sent for CLIA validation. Research results ascertained via samples #1 or #1A cannot be shared clinically until confirmed in a CLIA certified laboratory. **Abbreviations:** WES = whole exome sequencing, P/LP = pathogenic/likely pathogenic, CLIA = Clinical Laboratory Improvement Amendments to regulate laboratory based diagnostic testing.

#### Determination of a Gene List for Analysis and Return

In 2013, a team of genetics and genomics experts at Geisinger, in conjunction with a survey eliciting suggestions from national experts, developed a list of 76 genes for 27 conditions that builds upon the initial ACMG list of 56 genes. [3] The list includes 23 genes for cancer predisposition syndromes, 48 that confer risk for cardiovascular conditions, 2 for malignant hyperthermia, 2 for hereditary hemorrhagic telangiectasia and 1 for ornithine transcarbamylase deficiency (Additional File 1 Table 1). Participant consent to MyCode for receiving results is broad and allows flexibility to modify which results can be returned over time (Additional File 1 Figure 1). Additional types of genomic findings (e.g., pharmacogenomics) are under consideration for future analysis and return. Several overlapping candidate gene lists for risk identification have been developed by other groups.[3, 4, 8, 13] Many conditions on these lists share common characteristics such as under-recognition and undertreatment using standard clinical approaches, availability of confirmatory approaches to establish the medical diagnosis, evidence based preventive measures and/or treatments, and the possibility of a latency period before signs or symptoms develop.[3, 4]

#### Variant Analysis and Interpretation

To identify returnable results, sequence data is analyzed to determine those genetic variants associated with increased disease risk (i.e., expected pathogenic) and initiate confirmatory sequencing in a CLIA-certified environment. Only variants that are independently classified as pathogenic or likely pathogenic using established criteria[14] are confirmed by Sanger sequencing and reported to participants in the MyCode program. Variants of uncertain significance (VUS), although usually returned in diagnostic testing, are not returned to participants in this program. To minimize false positives for these conditions in an unselected population with an overall low prior probability, laboratory scientists perform a stringent analysis of variants that emphasizes specificity over sensitivity. Likely pathogenic results are treated as positive since this classification indicates a greater than 90% certainty that a variant is disease causing, while pathogenic classifications are associated with stronger evidence.[14] The absence of a confirmed result in a biobank participant is considered uninformative rather than negative due to limitations in the understanding of the impact of all variants (particularly those that are rare or unique) on disease and technical limitations of the sequencing process (e.g. difficulty detecting some deletions/duplications).

Genomic sequencing data generated for research use is stored electronically as variant call files (VCFs) that note differences between participants’ sequence data compared with a reference sequence. Files are attached to a study ID linked to individual participants. A GHS data broker manages these study IDs so that researchers cannot re-identify participants from VCFs. VCF data for the 76 genes - connected to the study ID, but absent of patient identifiers - are sent to a clinical laboratory (Partners HealthCare’s Laboratory for Molecular Medicine [LMM]) for analysis using their bioinformatics pipeline and variant interpretation process as described previously.[15] LMM utilizes stringent application of the ACMG criteria for variant classification in order to minimize clinical false positives.[14] When predicted pathogenic/likely pathogenic variants are identified in the VCF data, LMM requests DNA samples for these individuals from the biobank. The data broker holds the link to de-identified and identified samples and enables sample release, this time with all patient identifiers, for confirmation using an orthogonal technology (i.e., Sanger sequencing). An internal GHS pipeline for preliminary filtration of data to prioritize sequence files with suspected pathogenic or likely pathogenic variants has been developed and is being tested for performance with the goal to make this fully automated.

#### Participant Eligibility to Receive a Result

Participant eligibility to receive results is verified before CLIA confirmation of a suspected pathogenic or likely pathogenic result. Prior to requesting confirmation, the data broker verifies that individuals are alive and have not withdrawn from MyCode participation. Given the legacy of how the biobank was established, participants also must have completed an updated version of the consent that permits return of results and deposition in the electronic medical record. All individuals with an outdated consent are periodically contacted with a request to re-consent on a current consent that includes returning actionable results and placing those results in the medical record.

#### Confirmation and Reporting

Clinical laboratory reports are generated for CLIA confirmed findings and sent to the study team. This report is reviewed by the clinical team and uploaded into the participant’s EHR. A brief chart review is conducted to determine if the result was already known from prior genetic testing to tailor the result delivery process. In addition to the uploaded laboratory results report, participants and providers have access to user-friendly interpretive reports that have been developed at Geisinger. [16, 17] In the Geisinger MyCode Return of Results, the research budget covers the cost of confirmatory testing including the generation of the clinical laboratory report, and the initial discussion of the result with the patient (by a genetic counselor or geneticist and genetic counselor depending on the condition).

### 2. Delivery of Results to Healthcare Providers

Primary care providers (PCP) are key contributors in the results disclosure process. PCPs within GHS receive a message regarding patient results through the EHR 5-7 days before their patient is notified of the result (Additional File 1 Figure 2). Providers requested this pre-notification as they wanted to have some ownership of the return process whether they planned to return the result themselves, or work with the Genomic Medicine team to return the result. To provide support for providers about conditions they may not be familiar with, “genomic condition specific” 0.5 hour CME courses are available online for “just in time” education. [18] If a PCP does not wish to take an active role in disclosure or management of a genomic result, their patients are supported by the clinical genomics team. This process was developed through extensive consultation with system clinicians and clinical leadership. Assessment of physician experience with the return of results process is ongoing to allow iterative improvement.

A research coordinator notifies the PCP about patient results using a templated message in the EHR. The message includes the specific genomic result, the proposed plan for notifying the patient, and relevant available resources (Table 1). If review of the patient’s EHR identifies that the genetic finding was previously ascertained through clinical testing, this information is included in the message to the PCP. If a participant does not have a Geisinger PCP, then both the external PCP and the GHS provider with whom the patient met on the day of their consent are notified.

**Table 1.** Resources Developed for Participants and Providers

| Resource | Description |
| --- | --- |
| Provider Communication Following the Confirmation of a Research Result (Additional File 1 Figure 2) | Written notification to participant's PCP of confirmation of a research result. Includes laboratory report.<br>Notification is completed through the EHR for Geisinger providers, and with a letter for providers outside of Geisinger. |
| Patient Communication Following the Confirmation of a Research Result (Additional File 1 Figure 3) | Initial written notification to participant that a result exists and that they can call to find out more. Sent through the patient EHR portal or mailed letter. |
| Phone Script for The Disclosure of a Research Result (Additional File 1 Figure 4) | Script used by research assistant for an initial standardized phone conversation with patients to disclose the nature of the result and help direct next steps for follow-up |
| Patient Communication with Result Disclosure (Additional File 1 Figure 5) | Written notification detailing genomic result, importance of result sharing with family, and resource information. Sent after the patient has learned the nature of the result. |
| Unable to Contact Patient Letter (Additional File 1 Figure 6) | Written notification sent to patients who could not be reached by phone. Mailed using certified mail along with result-specific materials. |
| Laboratory Report with Result | Copy of the CLIA confirmed laboratory result |
|  | report for participant record keeping |
| Interpretive Test Reports | Patient and provider facing electronic reports that explain the genetic result and associated condition(s). Reports are gene-specific and interface with the EHR. |
| Patient Communication to Family Members for Result Sharing | Letter for participant to share with family members with information on result, importance of getting tested and how to obtain testing and care. A copy of the CLIA report is copied on the back. |
| Family Health History Questionnaire | Condition-specific family history questionnaire to gather family medical history and detect at-risk relatives |
| Patient Brief Condition Fact Sheet | Condition-specific information and resources for follow-up care, mailed to patients. |
| Provider Condition Specific Education Modules | Condition specific education modules for CME credit to inform providers available through a Geisinger educational platform to internal providers and through MedConcert for external providers. |
| Provider Clinical Guidance Sheet | A brief sheet with condition-specific information to facilitate care. |

Since one out of every three MyCode participants has a non-Geisinger PCP, it was important to determine how best to communicate this medically actionable information outside the GHS. An External Provider Liaison position was created to facilitate this communication. This liaison contacts PCPs outside of GHS by phone to notify them and mails a packet with a paper copy of the result as well as relevant information about the MyCode program and condition-specific resources. Additional providers are also notified when requested by the participant.

### 3. Disclosure of Results to Participants

#### Notifying Participants of Results

Participants are notified of results and provided with guidance for the appropriate next steps. The multi-step process includes an initial written notification, follow-up phone call, and a mailed information packet (Figure 2). This initial message, which notifies the participant that genomic testing has identified a finding important for their health, serves as a primer for return of results and encourages the participant to contact the MyCode team to learn more about the result (Additional File 1 Figure 3). It does not provide specific details about the result, and is either sent through the patient portal associated with the EHR or mail if the participant does not use the portal. Ten business days after the initial written notification, a research coordinator attempts to reach every participant who has not responded to the written notification. This phone disclosure is conducted using a result-specific script (Additional File 1 Figure 5) that highlights: 1. The nature of the result (i.e., that the result is associated with increased risk for certain diseases); 2. The result should be discussed with a healthcare provider; 3. The result should be shared with at-risk relatives; 4. A targeted family-history questionnaire should be completed; 5. A packet will be mailed with information about their result and letters to facilitate sharing the result with relatives (Additional File 1 Figure 5); and 6. They will receive a check-in call in 4 weeks. Participants are then asked to decide with which healthcare providers they would prefer to meet next (genomics, PCP, other specialist). Appointments with a genomics provider are available for scheduling at the time of telephone disclosure, during the 4-week follow-up call and by request.

**FIGURE 2:**
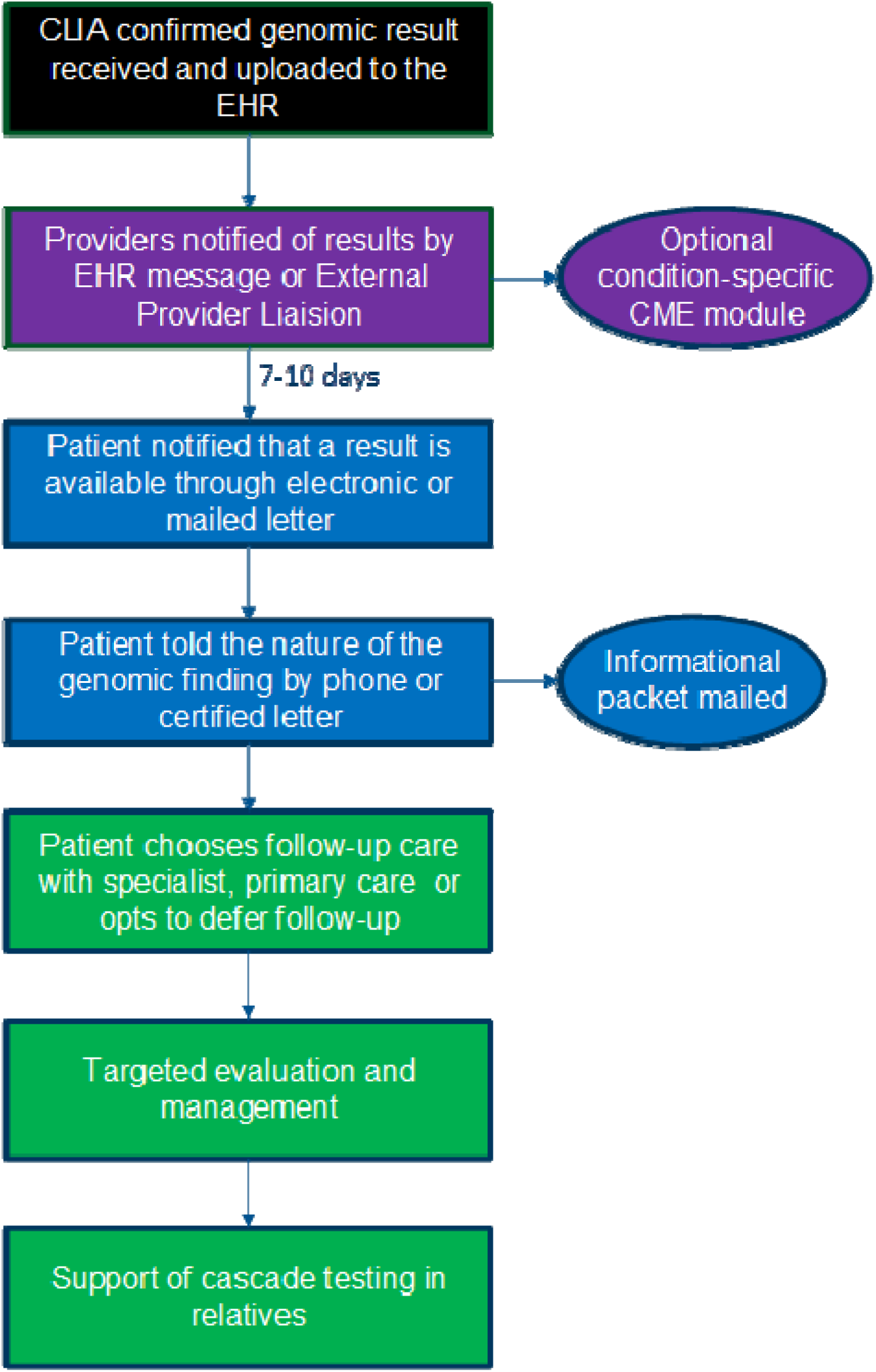
Figure 2 - Participant and Physician Notification Flow. The result enters the EHR (black) and then the process of notifying physicians begins (purple). Participants are notified of results through a process of multiple contact attempts (blue). Participants and their relatives receive care following their result disclosure (green).

The mailed information packet includes the genetic test report, information about the condition, letters for sharing their report with family members, information about completing a family history survey and details about how to schedule an appointment with genomics (Table 1). Participants who have not been reached by phone after 3 attempts are sent a packet by certified mail with the above materials and a letter indicating that we have been unable to contact them (6).

The recommendation for follow-up with a healthcare provider includes condition-specific information regarding targeted evaluation in either a specialty or primary care setting and genetic counseling. Appropriate specialists for each genomic condition have been identified throughout the system based on expertise and interest.

#### Genetics Assessment

Participants who choose to meet with the genomics team have the option to discuss their results in person or via phone or telemedicine. This genetic counseling session is often the first opportunity for an in-depth explanation of their genomic finding(s) and the associated medical conditions. Additional topics of discussion include: targeted review of personal and family histories, risks for variant-associated disease, condition-specific surveillance and management guidelines, inheritance patterns, and relatives with whom results should be shared. The genomics clinician facilitates guideline-based referrals for surveillance and management. Family history information is gathered to assess penetrance within the family (affecting risk assessment), to identify which relatives are at-risk, and to help understand psychosocially-relevant family dynamics. Participants have multiple avenues to provide their family history: an on-line family history tool, a paper family history form, or via a telephone call. Family history acquisition is targeted to genotype-associated phenotypes (i.e. clinical problems associated with the genetic variant).

Psychosocial support is provided throughout each session, with specific consideration of how the individual has been coping with the result (e.g., anxiety or distress), readiness to engage in recommended surveillance and management, and support for discussing the result with family members. Participants are encouraged to bring a family member for support.

### 4. Targeted clinical evaluations

Targeted clinical evaluations are crucial to the process of understanding genotype-phenotype correlations in this population. These targeted clinical evaluations may include assessment for relevant clinical manifestations, physical exam findings, pertinent past medical history, labs/diagnostics, detailed condition-specific family history, and consultation with specialists. Evaluations by appropriate clinical experts are encouraged, and are condition-specific. Targeted evaluations aid in identification of phenotypic findings that may require medical attention and inform research efforts to understand the natural history of the disease in this population genomics approach.

The genomic result is a risk marker, and not equivalent to a diagnosis (19). For patients whose clinical evaluation does not reveal a diagnosis of the associated genomic condition, the result is placed in the problem list of their EHR as a laboratory finding, to prompt appropriate ongoing follow-up. We have developed a framework in which participants are categorized into one of five diagnostic groups[19] based on whether the associated phenotype is present at time of evaluation.[18] The development of a care plan in this patient population is different from an indication-based evaluation, as it is a genetic finding that prompts the initial evaluation rather than the patient presenting with signs or symptoms of the disorder, or a family history. After the session, the evaluating clinicians work with the participant’s PCP to coordinate follow-up appointments with appropriate specialists and facilitate cascade testing of at-risk relatives.

### 5. Facilitating cascade testing of at-risk relatives

The genetic testing and counseling of at-risk relatives of participants who receive a result is critical for maximizing the population health impact of screening research exomes for actionable genomic results. Enough copies of the family letter are mailed to each participant to share with all living first-degree relatives, as ascertained by the research coordinator during the result disclosure phone call. During the week-four post disclosure check-in call, the research coordinator reminds the participant of the family letter, and asks if the result has been shared with any relatives. The importance of family sharing and follow-up is reiterated during this phone call. In addition, each genetic counseling session includes a focused discussion about the importance of sharing results with family members. Participants are reminded that their relatives may reach out to the genomics team with any questions and if desired, the team will assist with arrangements for testing. Family members who attend the initial genetic counseling appointment with the participant may pursue testing at that visit.

## RESULTS

As of July 3, 2017, 343 single gene results have been identified and returned using the methodology described above. These results were identified in 34 genes associated with increased risk for 18 conditions (Table 2). A publicly available list of the results that have been returned is updated monthly [18]. Return of results is in progress with certain conditions prioritized for return so the posted numbers should not be used to infer prevalence of the conditions in our population. In addition, not all patient samples have been formally reviewed in this ongoing project.

**Table 2.**
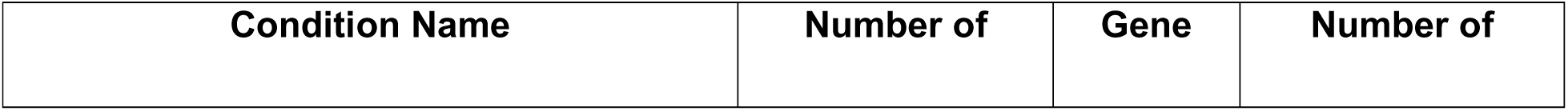
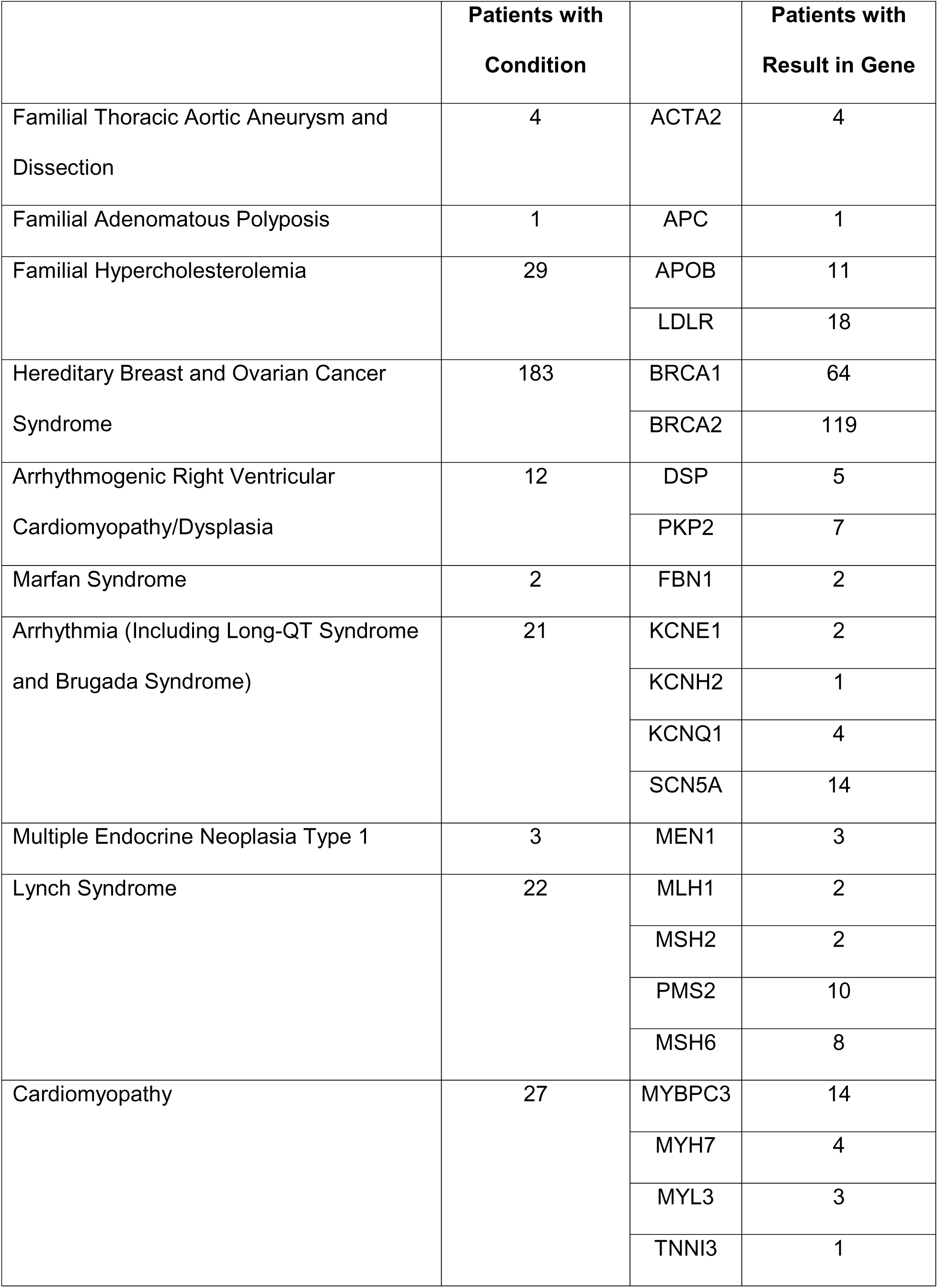
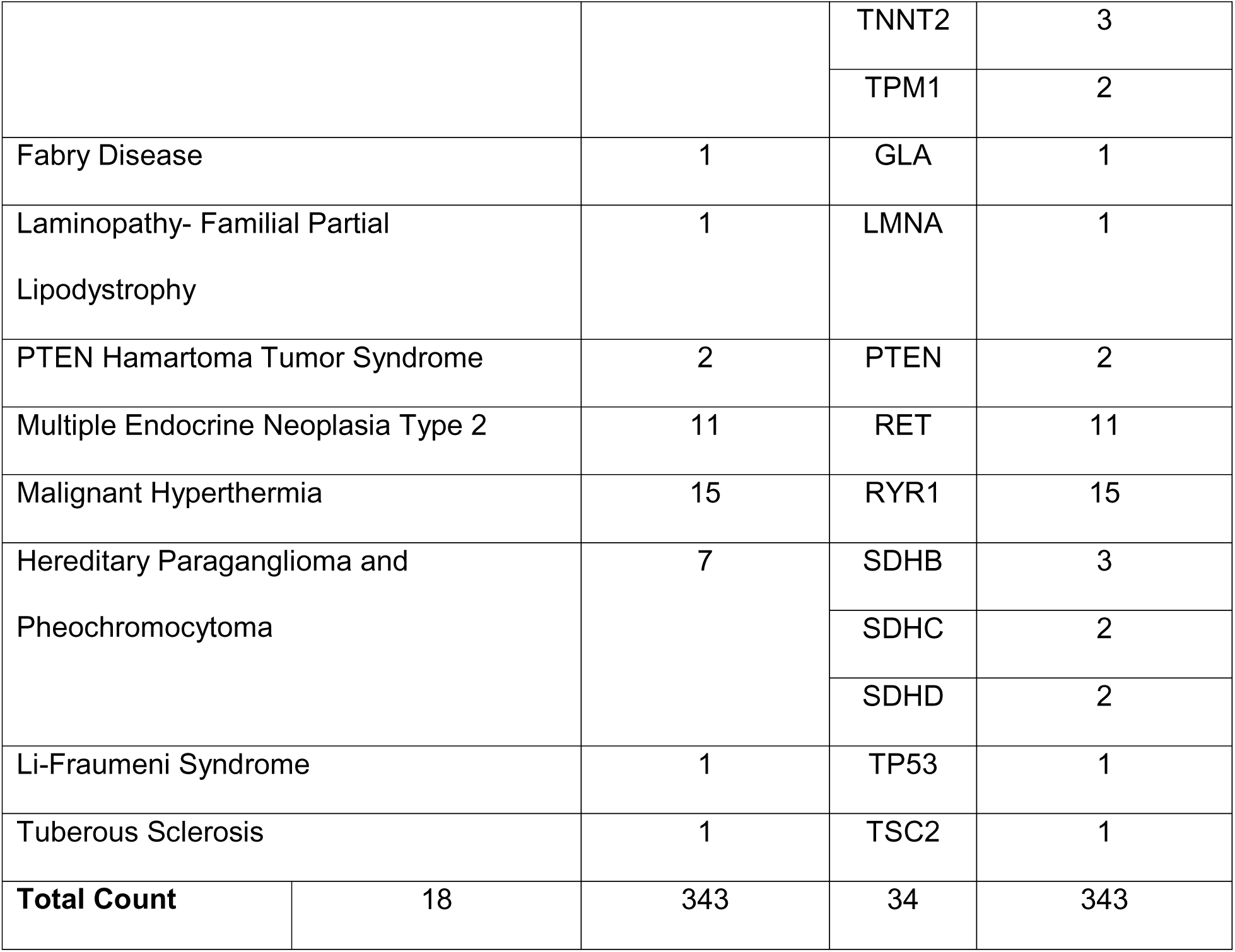
Results Delivered to Patients using this Genome-First Model as of July 3, 2017

| Condition Name | Number of | Gene | Number of |
| --- | --- | --- | --- |

|  | **Patients with**<br>**Condition** |  | **Patients with**<br>**Result in Gene** |
| --- | --- | --- | --- |
| Familial Thoracic Aortic Aneurysm and Dissection | 4 | ACTA2 | 4 |
| Familial Adenomatous Polyposis | 1 | APC | 1 |
| Familial Hypercholesterolemia | 29 | APOB | 11 |
|  |  | LDLR | 18 |
| Hereditary Breast and Ovarian Cancer Syndrome | 183 | BRCA1 | 64 |
|  |  | BRCA2 | 119 |
| Arrhythmogenic Right Ventricular Cardiomyopathy/Dysplasia | 12 | DSP | 5 |
|  |  | PKP2 | 7 |
| Marfan Syndrome | 2 | FBN1 | 2 |
| Arrhythmia (Including Long-QT Syndrome and Brugada Syndrome) | 21 | KCNE1 | 2 |
|  |  | KCNH2 | 1 |
|  |  | KCNQ1 | 4 |
|  |  | SCN5A | 14 |
| Multiple Endocrine Neoplasia Type 1 | 3 | MEN1 | 3 |
| Lynch Syndrome | 22 | MLH1 | 2 |
|  |  | MSH2 | 2 |
|  |  | PMS2 | 10 |
|  |  | MSH6 | 8 |
| Cardiomyopathy | 27 | MYBPC3 | 14 |
|  |  | MYH7 | 4 |
|  |  | MYL3 | 3 |
|  |  | TNNI3 | 1 |

|  |  |  |  |  |
| --- | --- | --- | --- | --- |
|  |  |  | TNNT2 | 3 |
|  |  |  | TPM1 | 2 |
| Fabry Disease |  | 1 | GLA | 1 |
| Laminopathy- Familial Partial Lipodystrophy |  | 1 | LMNA | 1 |
| PTEN Hamartoma Tumor Syndrome |  | 2 | PTEN | 2 |
| Multiple Endocrine Neoplasia Type 2 |  | 11 | RET | 11 |
| Malignant Hyperthermia |  | 15 | RYR1 | 15 |
| Hereditary Paraganglioma and Pheochromocytoma |  | 7 | SDHB | 3 |
|  |  |  | SDHC | 2 |
|  |  |  | SDHD | 2 |
| Li-Fraumeni Syndrome |  | 1 | TP53 | 1 |
| Tuberous Sclerosis |  | 1 | TSC2 | 1 |
| Total Count | 18 | 343 | 34 | 343 |

A total of 339 patients have received a result through this program, including 4 patients who have each received 2 distinct results (*BRCA2* and *APOB; BRCA1* and *LDLR; BRCA2* and *PKP2; SCN5A* and *LMNA*). One patient who received both an *SCN5A* and *LMNA* result had received only the *SCN5A* result in the initial result disclosure. The *LMNA* result linked to Familial Partial Lipodystrophy was identified by the lab, but reported as a research result to the team as the *LMNA* gene was on the return list for cardiomyopathy but not Familial Partial Lipodystrophy. In a consult with this patient, it became apparent that this individual had a known clinical diagnosis of Familial Partial Lipodystrophy and her healthcare providers worked with the laboratory with the verbal consent of the patient to have the report re-identified, issued to the patient, and placed in the EHR.

Among these 339 patients, 303 patients have had direct contact with the clinical genomics team as a part of the result disclosure process. Since this process is ongoing, 21 of the 339 patients are in the first phase of the return process where their providers have been notified, but the patients have not yet been notified of their results. A total of 15 patients have not been reached by the clinical genomics team and were sent the unable to contact patient packet as a certified letter (Additional File 1 Figure 6). One of these 15 patients has documentation in his chart of an interaction with primary care about his result. Overall, 304 out of 325 patients (93.5%) who have been contacted about their result are confirmed to have been successfully informed of their result using this process.

Additionally, 222 providers have been notified of patient results as a part of the result delivery process including 97 GHS primary care providers, 52 GHS specialists, and 73 external primary care providers.

## DISCUSSION

We describe a genome-first model for returning genomic results to unselected research participants within a health care system. The program integrates genomic data with electronic medical records to identify and return clinically actionable genomic variant information to participants and providers. The early stages of this program have been successful at reaching patients and their providers about results and demonstrate the feasibility of incorporating genomic sequence risk information into clinical care management. We are simultaneously building the evidence base to develop and continually refine clinical practice objectives for precision genomic medicine. This approach will allow for monitoring long-term outcomes related to health, health care utilization, psychosocial functioning, and economics. The foundation of the program focuses on variant identification, curation, and clinical interpretation. Although starting with a recognized list of actionable genes, the importance of the variant calling pipeline cannot be underestimated. Every effort is made to minimize the potential for harm and optimize the potential for improving health outcomes from the participant perspective. The implications of positive actionable findings in terms of potential significant and irreversible patient health decisions guided the process to ensure the most accurate variant classification possible.

Designing a program that met the needs of participants and providers required significant preparation and relied heavily on input from providers and participants. An important example of this was the request by providers involved instituting a delay in reporting results to patients, giving providers 5 days to investigate the result prior to releasing it to the patient. Participants endorsed the involvement of their providers and requested that results be returned in multiple formats (e.g. letter, phone, or in the patient EHR portal) given limited email or internet access. The clinical follow-up preferences of patients and their at-risk relatives is an area of future research interest extending from this project.

An important challenge of this program is that the participants who receive results in the early stages of the program will be managed using practices that often have been optimized to an population identified through traditional clinical diagnosis. This is particularly vexing given the uncertainty regarding disease penetrance in genotypically identified individuals.[20, 21] However, evidence generation in the context of the program, a key element of a LHS, will accrue knowledge that can then be applied to future participants with secondary genomic findings identified through clinical genomic sequencing—a problem that already exists and for which no evidence based recommendations exist.

Although the model applied here is specific to generating results from genomic screening, the same broad principles can be applied to other programs that search existing data for meaningful information. In particular, the EHR is a rich source of data with the potential to be mined to answer research and clinical questions.[22-24] The same basic steps of generating findings and ensuring they meet reasonable quality standards, communicating findings with clinicians and participants, and facilitating follow-up care would be applicable in this context.

Population based genomic screening programs have great potential to impact population health by enabling precision disease prevention and management, and can be optimized if implemented using principles of the LHS framework.[24] The model described here uses such a framework that can be adapted to other healthcare settings that aim to establish similar programs for returning genomic results to research participants. This program was developed using a LHS framework because the evidence base supporting the clinical use of genomic screening in a general population setting, while promising, remains immature. Use of the LHS framework to guide design, implementation, evaluation, and iteration enhances the likelihood that the program will improve health outcomes of importance to patients and providers and provide value to the healthcare system.

## List of Abbreviations

GHS: Geisinger Health System
MyCode: MyCode Community Health Initiative
ACMG: American College of Medical Genetics and Genomics
CLIA: Clinical Laboratory Improvement Amendment
EHR: Electronic Health Record
G76: Geisinger 76 Gene List
VUS: Variant of Uncertain Significance
VCF: Variant Call File
LMM: Partners Healthcare’s Laboratory for Molecular Medicine
PCP: Primary Care Provider
LHS: Learning Healthcare System
WES: Whole Exome Sequencing,
P/LP: Pathogenic/likely pathogenic

## Declarations

### Ethics approval and consent to participate

Approval for this study was received by the Geisinger Health System Institutional Review Board, Study number 2006-0258. Participants were consented in accordance with the approved protocol.

### Consent for publication

Not Applicable

### Availability of data and material

A list of the number and type of results returned for each condition through this program is updated monthly and is available on the GHS MyCode Website (https://www.geisinger.edu/en/research/departments-and-centers/genomic-medicine-institute/mycode-health-initiative) and is available upon request. Data on patient contact with our team is summarized within this article, our patient tracking dataset contains protected health information and is not available publicly.

### Funding

Robert Wood Johnson Foundation, Pioneer Award, Award #72832

Commonwealth of Pennsylvania, Department of Community and Economic Development, Award #C000058400

Frontier Communications

The Horace W. Goldsmith Foundation

Mericle Foundation

### Author’s contributions

All authors were involved in program development, design, implementation, and manuscript revision. MLBS, CZM, ALL, DML, MLGH and MFM were major contributors in drafting the manuscript. Data analysis for the results section was conducted by MLBS, ALL and LF. All authors read and approved the final manuscript.

## Acknowledgements

Not Applicable

## Additional File 1, Word Document contains the following

Additional File 1, Table 1: Existing Gene Lists for Return Of Findings Identified On Genome Sequencing

Additional File 1, Figure 1: Geisinger Mycode^®^ Community Health Initiative Consent Version 30

Additional File 1, Figure 2: Provider Communication Following the Confirmation of a Research Additional File 1, Figure 3: Initial Patient Communication Following the Confirmation of a Research Result

Additional File 1, Figure 4: Phone Script-Disclosure of A Research Result

Additional File 1, Figure 5: Post-Phone Disclosure Patient Letter

Additional File 1, Figure 6: Unable to Contact Patient Letter

